# Genome-wide identification of tomato xylem sap fitness factors for three plant-pathogenic *Ralstonia* species

**DOI:** 10.1101/2020.08.31.276741

**Authors:** Stratton J. Georgoulis, Katie E. Shalvarjian, Tyler C. Helmann, Corri D. Hamilton, Hans K. Carlson, Adam M. Deutschbauer, Tiffany M. Lowe-Power

## Abstract

Plant pathogenic *Ralstonia* spp. colonize plant xylem and cause wilt diseases on a broad range of host plants. To identify genes that promote growth of diverse *Ralstonia* strains in xylem sap from tomato plants, we performed genome-scale genetic screens (random barcoded transposon mutant sequencing screens; RB-TnSeq) in three strains spanning the genetic, geographical, and physiological range of plant pathogenic *Ralstonia: R. solanacearum* IBSBF1503, *R. pseudosolanacearum* GMI1000, and *R. syzygii* PSI07. Contrasting mutant fitness phenotypes in culture media versus in xylem sap suggest that *Ralstonia* strains are adapted to *ex vivo* xylem sap and that culture media impose foreign selective pressures. Although wild-type *Ralstonia* grew in sap and in rich medium with similar doubling times and to a similar carrying capacity, more genes were essential for growth in sap than in rich medium. Each strain required many genes associated with envelope remodeling and repair processes for full fitness in xylem sap. These genes were associated with peptidoglycan peptide formation (*murI*), secretion of periplasmic proteins (*tatC*), periplasmic protein folding (*dsbA*), synthesis of osmoregulated periplasmic glucans (*mdoGH*), and LPS biosynthesis. Mutants in four genes had strong, sap-specific fitness defects in all strain backgrounds: *murI, thiC, purU*, and a lipoprotein (RSc2007). Many amino acid biosynthesis genes were required for fitness in both minimal medium and xylem sap. Multiple mutants with insertions in virulence regulators had gains-of-fitness in culture media and neutral fitness in sap. Our genome-scale genetic screen identified *Ralstonia* fitness factors that promote growth in xylem sap, an ecologically relevant condition.

**Importance:** Traditional transposon mutagenesis genetic screens pioneered molecular plant pathology and identified core virulence traits like the type III secretion system. TnSeq approaches that leverage next-generation sequencing to rapidly quantify transposon mutant phenotypes are ushering in a new wave of biological discovery. Here we have adapted a genome-scale approach, random barcoded transposon mutant sequencing (RB-TnSeq), to discover fitness factors that promote growth of three related bacterial strains in a common niche, tomato xylem sap. Fitness of wild-type and mutants show that *Ralstonia* spp. are adapted to grow well in xylem sap from their natural host plant, tomato. Our screen identified multiple sap-specific fitness factors with roles in maintaining the bacterial envelope. These factors include putative adaptations to resist plant defenses that may include antimicrobial proteins and specialized metabolites that damage bacterial membranes.

## Introduction

*Ralstonia solanacearum, R. pseudosolanacearum*, and *R. syzygii* (hereafter, “*Ralstonia*”) comprise the monophyletic but diverse species complex of plant wilt pathogens (1, 2). *Ralstonia* strains are adapted to plant hosts belonging to over 50 botanical families and are distributed worldwide in in warm tropics and temperate subtropical highlands with year-round precipitation (3–7). Most *Ralstonia* strains invade plant roots, gain entry to the water-transporting xylem vasculature, and spread systemically, which disrupts xylem function and fatally wilts the host plant (8).

Key molecular pathogenesis, virulence, and fitness traits have been identified by studying model strains like *R. pseudosolanacearum* GMI1000. However, comparative genomics shows that the *Ralstonia* species complex is genetically heterogeneous (1, 9–11). Average nucleotide identity (ANI) between strains in the different species ranges between 90.8-92.7% (1), which is less than the 95% ANI threshold widely used for species borders. An average *Ralstonia* genome contains approximately 5,000 genes, but approximately 2,900 core gene families are core to the *R. solanacearum* species complex (unpublished data from Vinatzer and Lowe-Power labs).

Accordingly, ecological niches, host ranges, and physiological traits vary widely among *Ralstonia* strains (1, 12–14). Developing and studying new model strains (15) that capture the phylogenetic diversity of the *Ralstonia* species complex will provide a more complete view of the biology of these globally important pathogens.

We hypothesized that genes in diverse *Ralstonia* strains contribute to their growth in xylem sap of tomato plants, a common host. In this study, we used a RB-TnSeq *mariner* transposon library with over 10^8^ possible barcoded Tns (16) to create genome-wide transposon insertion mutant libraries of three tomato-pathogenic *Ralstonia* strains: *R. pseudosolanacearum* GMI1000, *R. solanacearum* IBSBF1503, and *R. syzygii* PSI07. We mapped the transposon insertion sites and predicted core essential genes. We performed genetic screens to identify fitness factors that influence competitive growth of strains in *ex vivo* xylem sap from healthy susceptible tomato, as well as minimal and rich culture media. Comparison of genes’ fitness contributions between conditions revealed sap-specific fitness factors and genes that contribute to fitness in both sap and in culture media.

## Results

### Adapting RB-TnSeq for plant pathogenic *Ralstonia* isolates

To identify genes required for *Ralstonia* fitness, we constructed barcoded transposon insertion mutant libraries in three *Ralstonia* species complex backgrounds: *R. pseudosolanacearum* GMI1000, *R. solanacearum* IBSBF1503, and *R. syzygii* PSI07. We selected these isolates because they are all virulent on tomato and because there are high-quality, complete genomes available for each strain. These diverse strains share 76-83% of their gene content with average nucleotide identities of 91-93% (1), which we hypothesized would allow us to identify strain-specific and conserved fitness factors required by the species complex.

The mutant libraries were created by conjugation with the pKMW3 *E. coli* donor library (16), which carries a pool of over 10^8^ *mariner* transposon delivery vectors in which each transposon is marked by a unique 20 base pair sequence. These unique sequences function as DNA “barcodes”. After conjugation, we selected and pooled approximately 10^5^ transposon insertion mutants per strain library and mapped the insertion site of each barcoded transposon. Table 1 shows detailed summary statistics of each mutant library. For barcode sequencing (BarSeq) mutant fitness assays, we calculate fitness contributions using transposons that had inserted into the central 80% of the open reading frame (ORF) because insertions at a gene’s N- and C-termini are less likely to impair protein function.

**Table 1:** Properties of barcoded *mariner* mutant libraries.

| Mutant Library | Total genes in genome | Genes with Fit scores: # (% total) | Library Size<sup>a</sup>: # mutants | Mutants per gene in fitness calculations: Median (Mean) | Estimated Essential Genes<sup>b</sup>: # (%) |
| --- | --- | --- | --- | --- | --- |
| **GMI1000** | 5,204 | 4,003 (77%) | 90,016 | 11 (17.1) | 456-470 (8.7-9.0%) |
| **PSI07** | 4,968 | 4,057 (82%) | 190,155 | 22 (35.7) | 454-487 (9.5-9.8%) |
| **IBSBF1503** | 4,897 | 3,809 (78%) | 104,510 | 12 (20.4) | 454-487 (9.3-10.0%) |

<sup>a</sup>Measured by # of DNA barcodes mapped to a specific insertion site.
<sup>b</sup>Two pipelines (described in Methods) were used to predict gene essentiality based on the distribution of transposon insertions in open reading frames.

The barcoded mutant libraries cover 79-86% of the protein coding genes of each strain. There are several reasons why a given gene may be unrepresented in the mutant libraries. The *mariner* transposon we used inserts into TA dinucleotide sites. Strains GMI1000, IBSBF1503, and PSI07 have 48, 42, and 101 genes that could not be mutagenized because they lack TA sites. Table S1 lists genes that lack fitness data and the number of TA sites in the central 80% of each gene. Additionally, it is not possible to uniquely map transposon insertions in repetitive, indistinguishable genomic regions like the 31 kb tandem duplication in the GMI1000 genome. Furthermore, if these genes contribute to fitness in a condition, they would be functionally redundant with their identical paralogs, mitigating any potential impact on fitness. Finally, a subset of the genes without transposon insertions are likely essential or near-essential for growth in rich medium, resulting in their failure to grow during library construction and exclusion from our libraries.

### Predicting genes essential for *Ralstonia* growth in rich medium

We predicted essentiality of genes in each strain (Table S1) using the Bio::TraDIS pipeline (17) and our previously described analysis pipeline (18). Briefly, both methods predict gene essentiality based on genes that lacked transposon insertions in the central 80% of the coding region. After setting cut-offs to exclude short genes (“Above length cutoff” in Table S1), which are more likely to lack transposon insertions, 450-500 genes were predicted to be essential per strain by at least one pipeline, while 393-396 genes per strain were predicted to be essential by both pipelines (Table S1). To compare results between strains, we identified orthologous genes in GMI1000, PSI07, and IBSBF1503 using the JGI-IMG (19) Genome Gene Best Homologs tool. A core set of 244 genes was predicted to be essential by both pipelines in all three strains (Fig S1) (20). We inspected the predicted essential gene set for a positive control, the *speC* ornithine decarboxylase. We previously showed that a GMI1000 *speC* mutant (RSc2365) is a putrescine auxotroph that cannot not grow in rich medium unless it is supplemented with putrescine (8); both pipelines predicted *speC* was essential in GMI1000 and PSI07. Bio::TraDIS, but not our own pipeline, predicted *speC* to be essential in IBSBF1503.

To visualize which cellular processes are predicted essential for growth in rich medium, we used KEGG Mapper (21) to query the KEGG database with the GMI1000 locus tags of the 244 genes predicted to be essential in all three strains. As expected, many of these genes are involved in central cellular functions, such as DNA replication, transcription, translation, mRNA degradation, and lipid and peptidoglycan biosynthesis. Several genes from central carbon metabolism and respiration are predicted to be essential, including components of the Entner-Doudoroff pathway, the pentose phosphate pathway, the TCA cycle, NADH dehydrogenase, and ATP synthase. Many genes involved in cofactor and vitamin biosynthesis (ubiquinone, coenzyme A, thiamine, heme, and folate) are predicted to be essential. Genes that encode biosynthetic steps of several amino acids, including threonine, arginine, lysine, and histidine, also appear to be essential for growth in rich medium.

### Growth of *Ralstonia* species representatives in xylem sap

To design an RB-TnSeq experiment with a sufficient number of cell doublings to quantify mutant fitness defects, we empirically determined the carrying capacity and doubling rate of *Ralstonia* in xylem sap. We compared growth of *R. pseudosolanacearum* GMI1000 in rich medium, minimal medium, or xylem sap harvested from greenhouse-grown Moneymaker tomato (Fig S2A-B). The carrying capacities of xylem sap and rich medium were higher than the carrying capacity of minimal medium: 3.1-3.5 × 10^9^ CFU/ml in sap, 2.4-2.6 × 10^9^ CFU/ml in rich medium, and 2.2-4.6 × 10^8^ CFU/ml in minimal medium. Both xylem sap and rich medium cultures exceeded 1 × 10^8^ CFU within 24 hours, while minimal medium cultures had an extended lag time. We were surprised by the high carrying capacity supported by xylem sap based on our previous experience growing diverse *Ralstonia* in xylem sap from a different susceptible tomato (Bonny Best) grown in environmentally controlled growth chambers (8, 22). To determine whether the growing conditions or plant cultivar influenced the saps’ carrying capacity, we collaborated with researchers at University of Wisconsin—Madison to repeat this experiment with sap from growth chamber-grown Moneymaker and Bonny Best tomato in all three of our study strains. Each *Ralstonia* isolate grew to a higher carrying capacity in xylem sap from Moneymaker plants (3.3-6.6 × 10^9^ CFU/ml) than sap from Bonny Best plants (3.6 × 10^7^-5.5 × 10^8^ CFU/ml) (Fig S2C-E).

To estimate the doubling rate in xylem sap and rich and minimal culture media, we compared growth of GMI1000, PSI07, and IBSBF1503 with frequent sampling intervals (every 4 hours, Fig 1A). Strains grew slowest in minimal medium and the fastest in xylem sap (Fig 1B). The maximum doubling rate in xylem sap was 120-131% higher than in rich medium (P<0.005 by ANOVA with Tukey’s multiple comparison’s test). The higher growth rate and carrying capacity in xylem sap compared to rich medium indicate that *Ralstonia* strains GMI1000, IBSBF1503, and PSI07 are well-adapted to growth in xylem sap from Bonny Best tomato plants.

**Fig 1.**
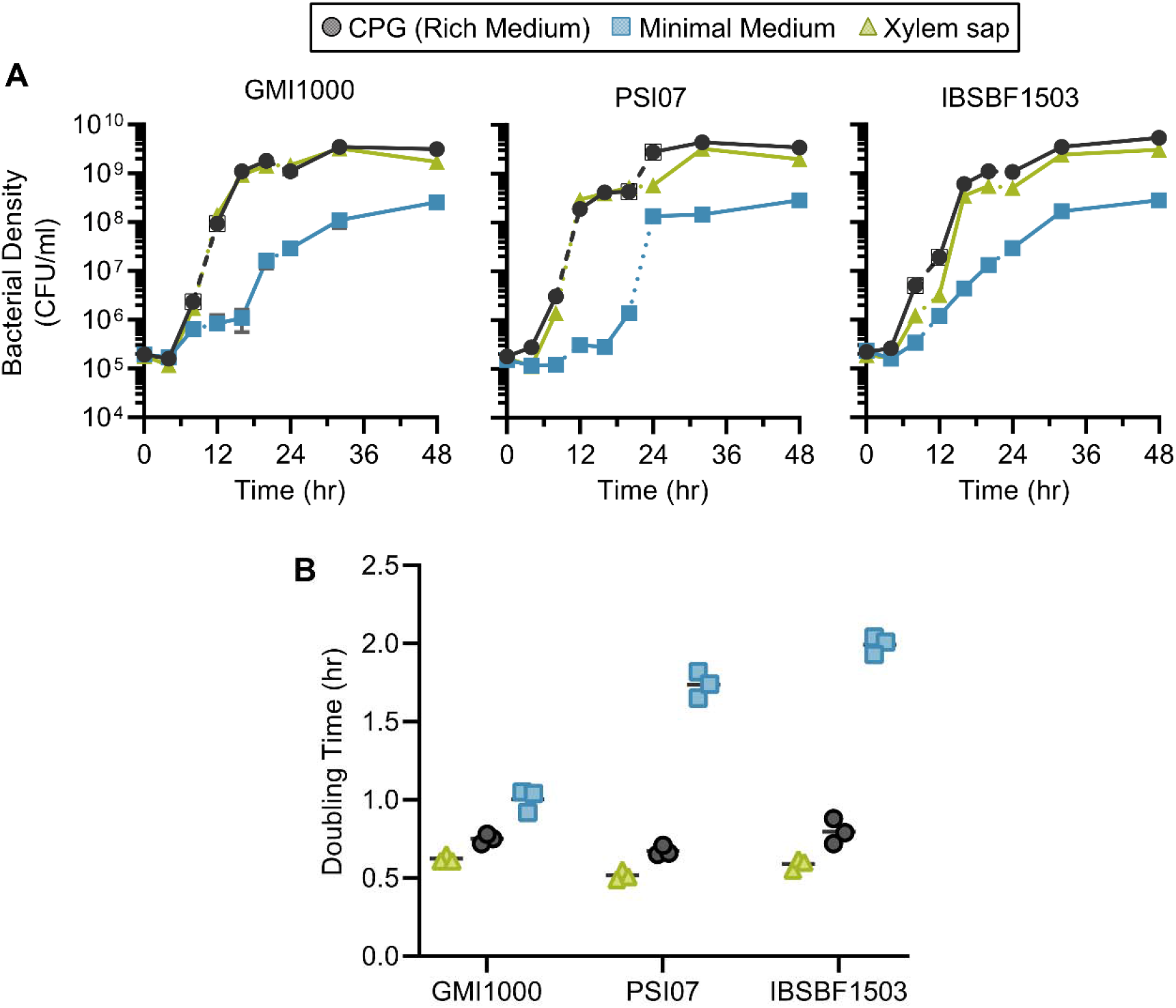
Growth of *Ralstonia* strains in xylem sap is similar to growth in rich medium. *Ralstonia* strain GMI1000, strain PSI07, or strain IBSBF1503 were grown in rich medium (CPG), quarter-strength M63 minimal medium (MM) or xylem sap from Moneymaker tomato plants. (**A**) Strains were inoculated at 10^5^ CFU/ml into CPG, MM, or xylem sap. All cultures were set up with 3 biological replicates (individual overnight cultures and individual batches of xylem sap) and were performed in technical duplicate. Solid lines indicate time points sampled from the same culture; the 12-24 hour timepoints were from an independent culture, but 0-8 and 24-48 hour timepoints were taken from the same culture. Error bars indicate standard deviation. (**B**) The minimum doubling time between contiguous 4 hour measurements was calculated per biological replicate. Lines indicate mean. Per strain, the doubling times in all conditions were significantly different from each other (adjusted P<0.05 by ANOVA with Tukey’s multiple comparisons).

### RB-TnSeq experiments identify mutants with altered fitness in xylem sap and minimal medium

We used RB-TnSeq to identify genes that influence fitness in *ex vivo* xylem sap, minimal medium, and rich medium. Mutant abundance before and after selective competitive growth was measured by BarSeq. In BarSeq, each gene is assigned a fitness score (“Fit” score), which is a weighted average of the log_2_ ratios of each barcoded mutant’s final abundance divided by its abundance before growth in the selective condition (the “time 0” abundance) (16). In total, our BarSeq assays measured fitness contributions for approximately 4,000 genes per strain. Multiple independent transposon mutants in the same gene are used to calculate each gene’s Fit score (Table 1). We calculated Fit scores and t-like test statistics for each gene in each condition (Table S2). Fitness data are also publicly available on the interactive Fitness Browser (http://fit.genomics.lbl.gov/). Because Fit scores are on a log_2_ scale, a score of −5 shows in principle that those mutants replicated for 5 fewer generations than the population. In practice, we found that mutants with the strongest fitness defects had Fit scores that exceeded the number of generations that we measured by dilution plating. For example, GMI1000 *murI* (RSc1956) had a mean Fit score of –8.6 in xylem sap even though the population grew for only 7 generations. However, most Fit scores were in the expected ranges for each TnSeq trial.

There were trends in the number genes associated with gain or loss of fitness between growth conditions (Fig 2A-C). There were more mutants with fitness defects in xylem sap (n=37-55) and minimal medium (n=28-42) than in rich medium (n=8-28) (Fit<-1; t-like statistic <-2.5 in all trials). Logically, mutants with strong fitness defects in rich medium are excluded from the libraries because they would not have grown on the selection plates. Interestingly, we observed that more mutants had gain-of-fitness phenotypes in minimal (4–20) and rich media (2–19) than in xylem sap (0-1). We next determined which genes influence fitness in a single condition and which genes have intersecting fitness contributions in two or more conditions (Fig 2D). The intersection analysis reveals that each strain has many genes that contribute to fitness in xylem sap only, in minimal medium only, or in both xylem sap and minimal medium.

**Fig 2.**
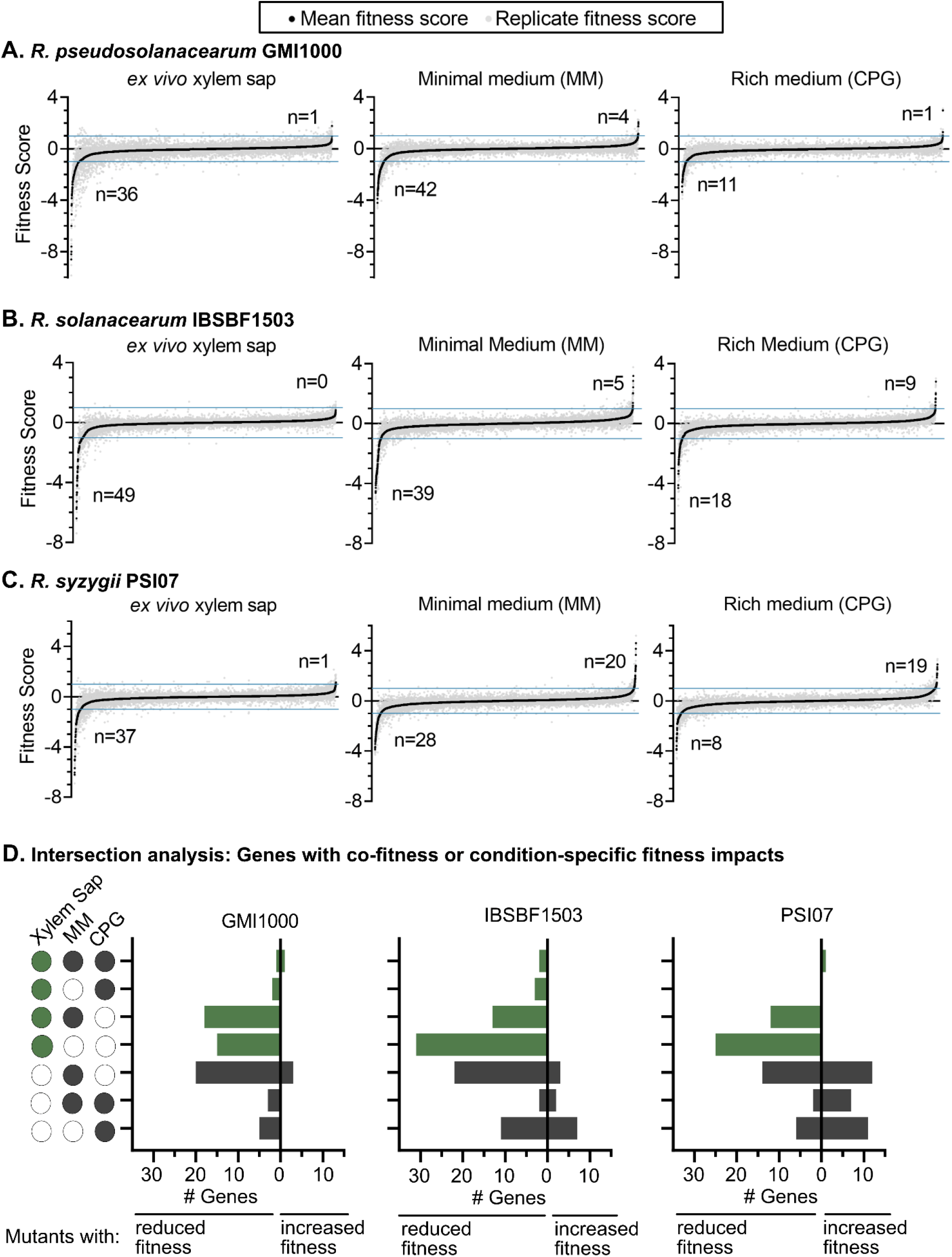
Summary of RB-TnSeq data after growth of mutant libraries in xylem sap, minimal medium (MM), or rich medium (CPG). (A-C) Ranked Fit scores for all measured genes in CPG, MM, and xylem sap for (A) *R. pseudosolanacearum* GMI1000, and (B) *R. solanacearum* IBSBF1503, and (C) *R. syzygii* PSI07. The number of genes that influence fitness in each condition is indicated (average Fit > 1 or <-1; t-like test statistic >|2.5| all three RB-TnSeq trials). (D) Intersection analysis quantifying number of genes that influence fitness in a single condition or that have co-fitness in multiple conditions. Green bars indicate genes that influence fitness in sap and grey bars indicate genes that only influence fitness in culture media. The intersection visualization is modeled on UpSet Plots (66), an alternative to Venn Diagrams.

### *Ralstonia* require multiple envelope-associated genes for fitness in xylem sap

Our intersection analysis identified envelope-associated genes as fitness factors in xylem sap in all three strains, in addition to other strain-specific xylem sap fitness factors (Fig 3). The strongest xylem sap fitness factor in all strains was the *murI* gene (RSc1956) with a Fit score of < −5 in all strains. MurI is a glutamate racemase that converts L-Glutamate to D-Glutamate, which is incorporated into peptide cross-linkages in peptidoglycan. The *murI* gene is required for multiple *Ralstonia* strains to grow in tomato stems (23, 24). For all three study strains, other xylem sap fitness factors included *tatC* (involved in Tat-dependent protein secretion), *dsbA* which catalyzes disulfide bonds in periplasmic proteins, and a lipoprotein of unknown function (RSc2940, RSc0285, and RSc2007, respectively). BLAST analysis reveals that this lipoprotein is present in genomes of *Ralstonia, Cupriavidus, Paraburkholderia, Burkholderia*, which are all genera in the *Burkholderiaceae* family.

**Fig 3.**
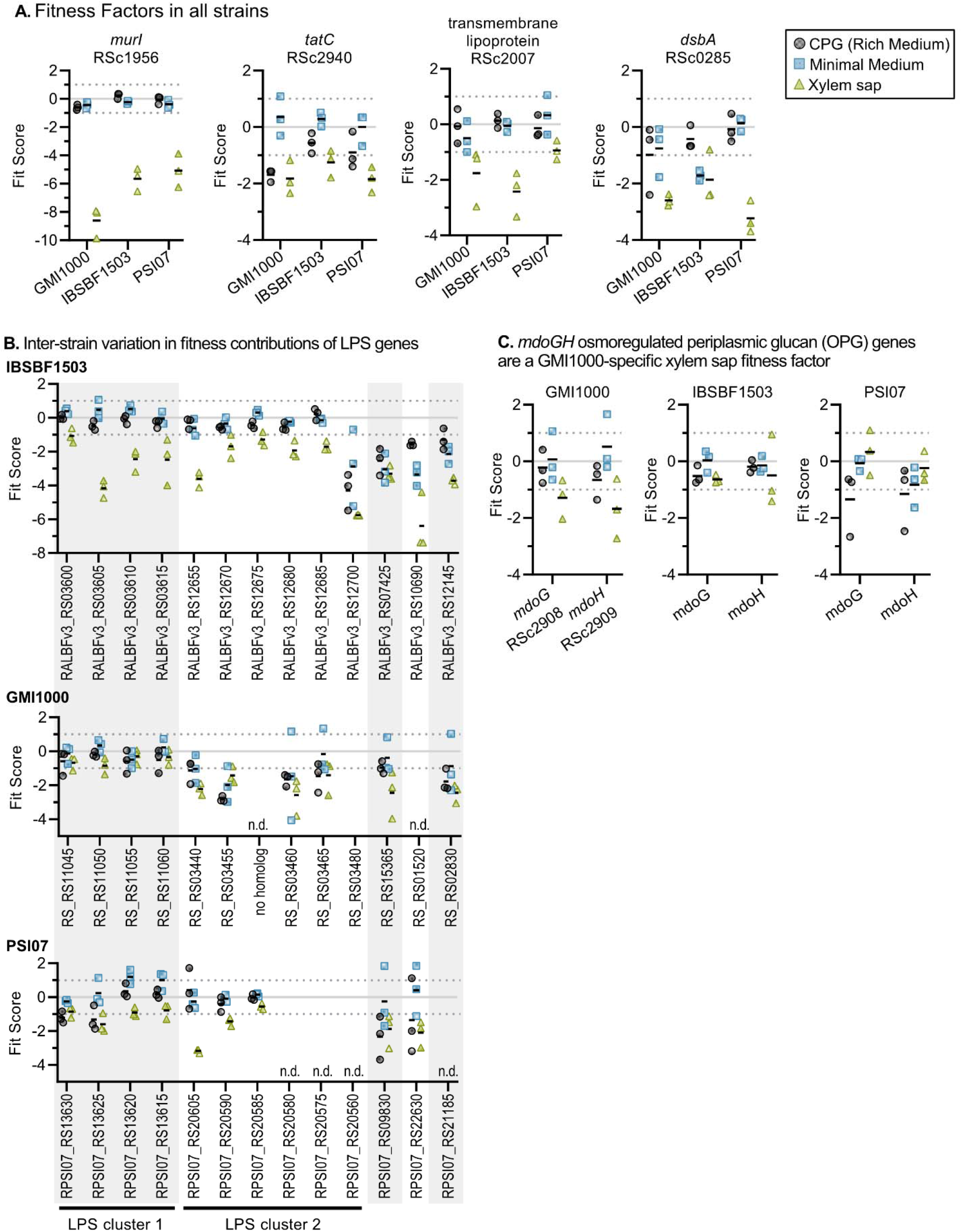
Envelope-associated genes contribute to fitness in xylem sap. Fit Scores of xylem sap fitness factors in one or more *Ralstonia* strains. Graphs show fitness contributions of (A) Four envelope-associated genes that are sap fitness factors in all three strains, (B) LPS genes that are xylem sap fitness factors for at least one strain, and (C) periplasmic glucan biosynthesis genes that are xylem sap fitness factors for GMI1000. Graphs show Fit scores of n=3 TnSeq trials and bars show the mean. Full Fit scores and t-like statistics are shown in Table S2.

IBSBF1503 requires 13 different LPS genes for full fitness in xylem sap, and several of these genes are important for fitness in xylem sap in PSI07 and GMI1000 as well (Fig 3B). Though conserved in all three strains, LPS genes colocalized in LPS cluster 1 had stronger fitness phenotypes in IBSBF1503 (−4.2 < Fit < −1.1) than in GMI1000 (−0.9 < Fit < −0.3) and PSI07 (−1.6 < Fit < −0.8). These results corroborate previous work in *Ralstonia* strain Pss4 indicating that mutants with structural defects in LPS also have reduced growth *in planta* (25) while highlighting subtle interstrain differences in the exact orthologs involved.

Other strain-specific envelope proteins identified in our analysis include the finding that GMI1000 requires *mdoGH* (RSc2908 and RSc2909) during growth in xylem sap (Fig 3C), which putatively encode enzymes that synthesize periplasmic branched oligoglucans. Since periplasmic oligoglucans can act as osmoprotectants to Gram-negative bacteria (26), and *mdoGH* contributes to the growth of several plant pathogens the host (27), these enzymes may play a role in protecting GMI1000 from environmental stress posed during growth in xylem sap.

### Metabolic functions required for full fitness in xylem sap

Many of the xylem sap fitness factor genes are associated with bacterial metabolism. Consistent with previous studies that showed *Ralstonia scrA* sucrose catabolism mutants (RSp1285) have growth defects in tomato stems (28, 29), *scrB* (RSp1284) sucrose catabolism mutants also had reduced fitness in xylem sap in our TnSeq experiments. The fitness defect of *scrB* mutants was stronger in the IBSBF1503 background than in the GMI1000 or PSI07 backgrounds (Fit = −2.0, −1.3, and −0.9, respectively). Overall, our findings validate that sucrose is an important carbon source in xylem sap although *Ralstonia* can catabolize other carbon sources present in sap (8, 30, 31).

All strains required amino acid biosynthesis genes for fitness in xylem sap and minimal medium (Fig 4). Mutants lacking Ser, Cys, Met, Phe, and Trp biosynthesis genes had fitness defects in both xylem sap and minimal medium. Although all strains required several Glu, Pro, and branched chain AA (Leu/Ile) biosynthesis genes for growth in minimal medium, only GMI1000 consistently required these genes for full growth in xylem sap. Our TnSeq results are consistent with xylem sap metabolomic studies that indicate that multiple tomato cultivars lack Cys and Trp in sap (8, 31) Although Ser, Met, and Phe are all present in xylem sap, our TnSeq results indicate that *Ralstonia* requires many *de novo* amino acid synthesis enzymes for full fitness in xylem sap.

**Fig 4.**
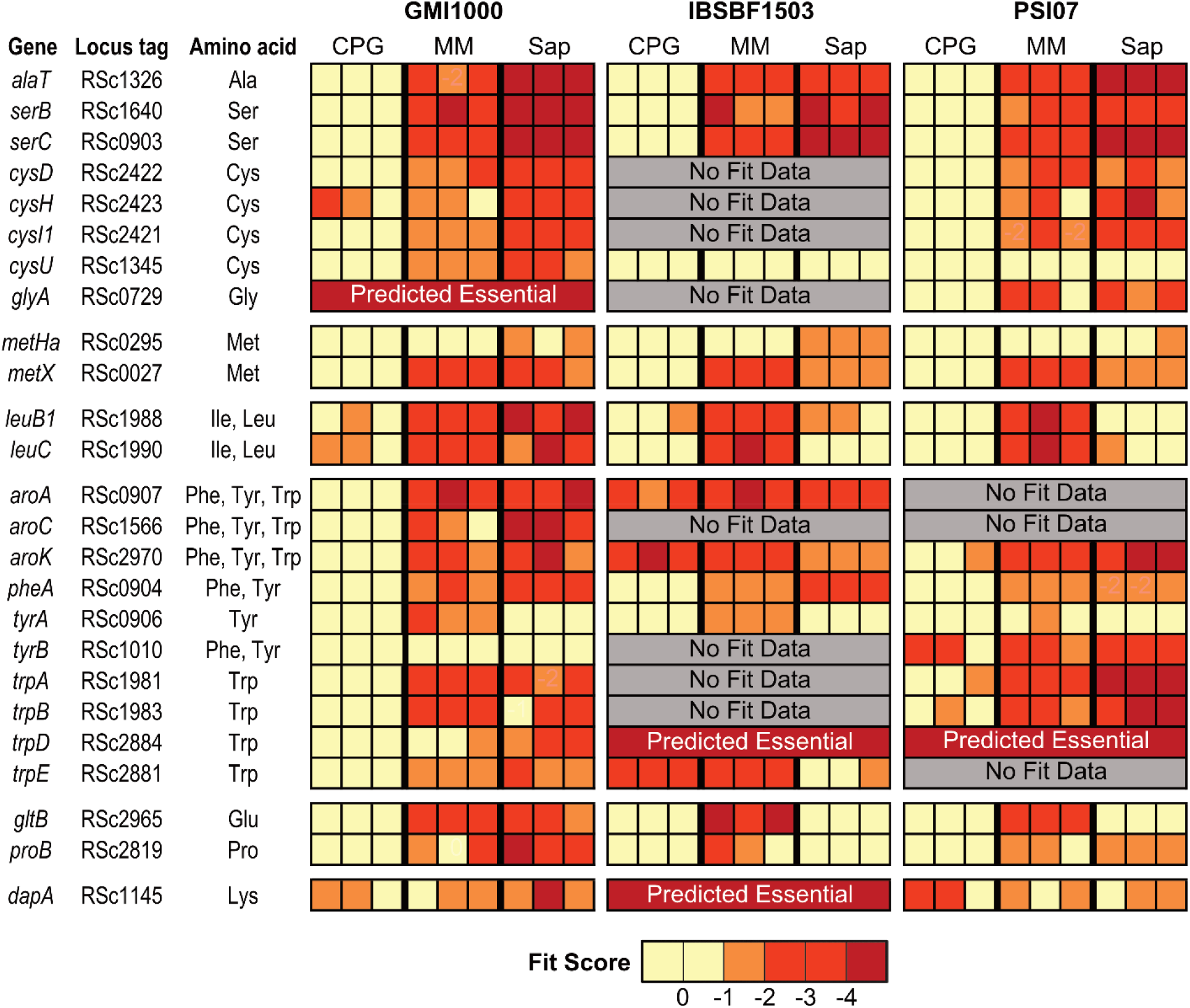
Amino acid auxotroph mutants have fitness defects in minimal medium (MM) and/or xylem sap. Fit scores of amino acid biosynthesis mutants with reduced fitness in sap, MM, or both. Gene name and locus tag for the GMI1000 gene are displayed. Full Fit scores and t-like test statistics are shown in Table S2.

Multiple cofactor biosynthesis mutants had xylem sap fitness defects in one or more strains (Fig 5). All three strains required the thiamine biosynthesis gene *thiC* for full growth in xylem sap (Fit −1.0 to −2.4). Similarly, all strains strongly required *purU* (RSc1873) for growth in xylem sap (Fit −3.9 and −6.0), with weaker fitness defects in minimal medium (Fit −0.3 to −1.4). PurU is a putative formyltetrahydrofolate deformylase that hydrolyzes 10-formyltetrahydrofolate into formate and tetrahydrofolatic acid (THFA). Because THFA is a cofactor for biosynthesis of many amino acids and nucleic acids, it is possible that PurU is necessary for *Ralstonia* to recycle THFA in order to synthesize amino acids. Interestingly, only one of the three strains required cobalamin biosynthesis to grow in xylem sap. Eighteen IBSBF1503 cobalamin biosynthesis mutants had subtle but consistent fitness defects in xylem sap with Fit scores that ranged −0.9 to −1.4 (Fig 5B).

**Fig 5.**
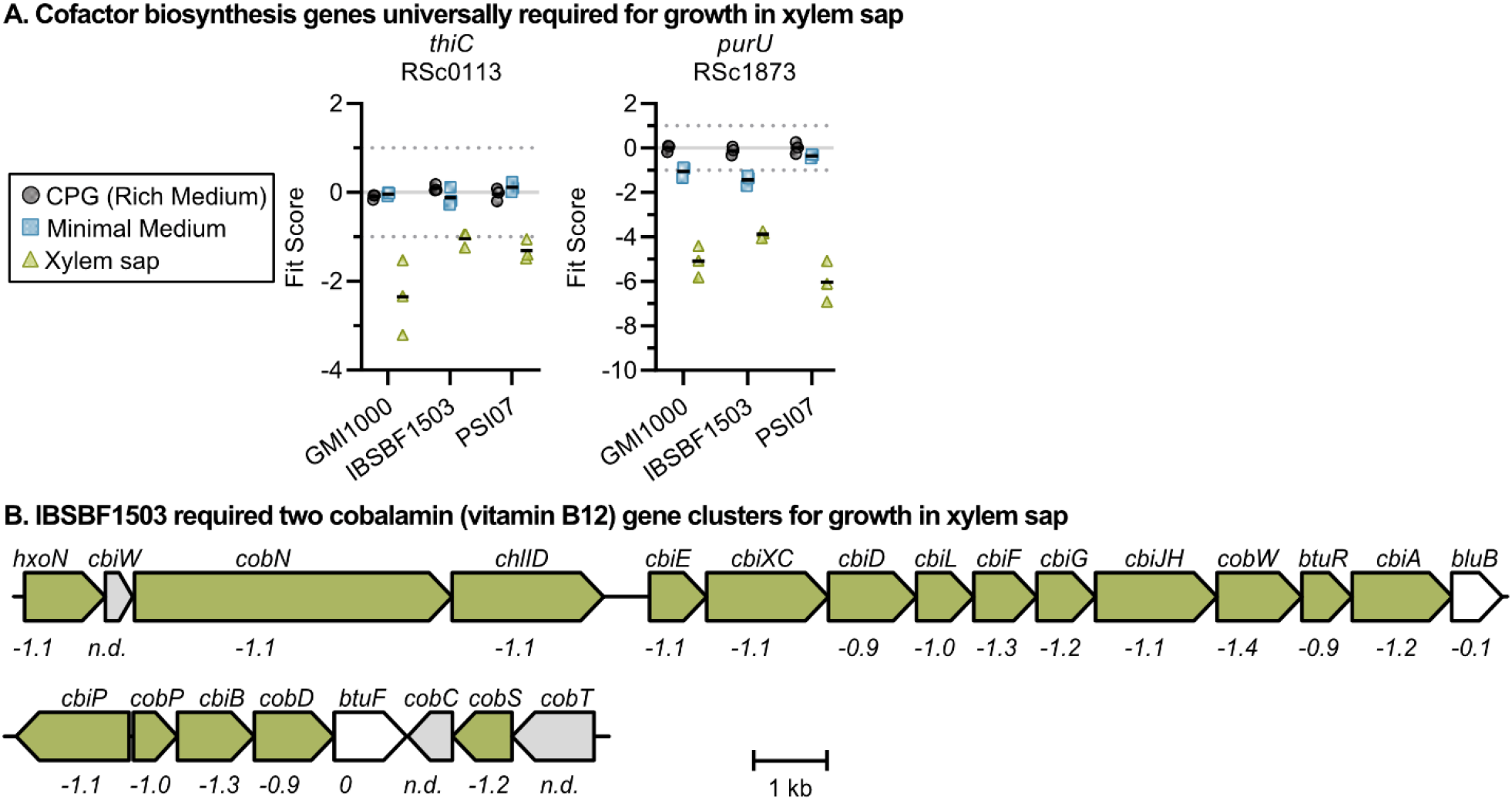
Mutants lacking cofactor biosynthesis genes have fitness defects in xylem sap. (A) A thiamine and tetrahydrofolic acid gene were both xylem sap fitness factors in all three strains. Graphs show Fit scores of n=3 TnSeq trials and bars show the mean. (B) *Ralstonia* IBSBF1503 required cobalamin biosynthesis for growth in xylem sap. The xylem sap fit scores of each gene are written below the genes. “n.d.” indicates fitness was not determined because mutants were not present in the RB-TnSeq library. Full Fit scores and t-like test statistics are shown in Table S2.

### Mutants with improved fitness

Many mutants with gains-of-fitness phenotypes had transposon insertions in regulatory genes (Fig 6). Consistent with previous results showing that the PhcA quorum sensing regulator mediates a trade-off between fast growth at low cell density and expression of *in planta* fitness traits at high cell density (32, 33), GMI1000 and IBSBF1503 *phcA* mutants had improved growth in all tested conditions (Fig 6A). Unexpectedly, PSI07 *phcA* mutants behaved differently. Although PSI07 *phcA* mutants had increased fitness in minimal medium (Fit: +1.9), they had strongly reduced growth in xylem sap (Fit: −2.8). Contrasting phenotypes of *phcA* mutants in different wildtype backgrounds highlights the genetic and regulatory diversity within plant-pathogenic *Ralstonia*. At high cell densities, PhcA positively regulates production of an energetically costly extracellular polysaccharide (EPS) among other traits (33, 34). Transposon insertion in other regulatory genes that activate expression of *eps* and other virulence genes (*phcBSR, vsrAD*, and *rpoS*) (35, 36) also increased fitness in one or more strains (Fig 6A).

**Fig 6.**
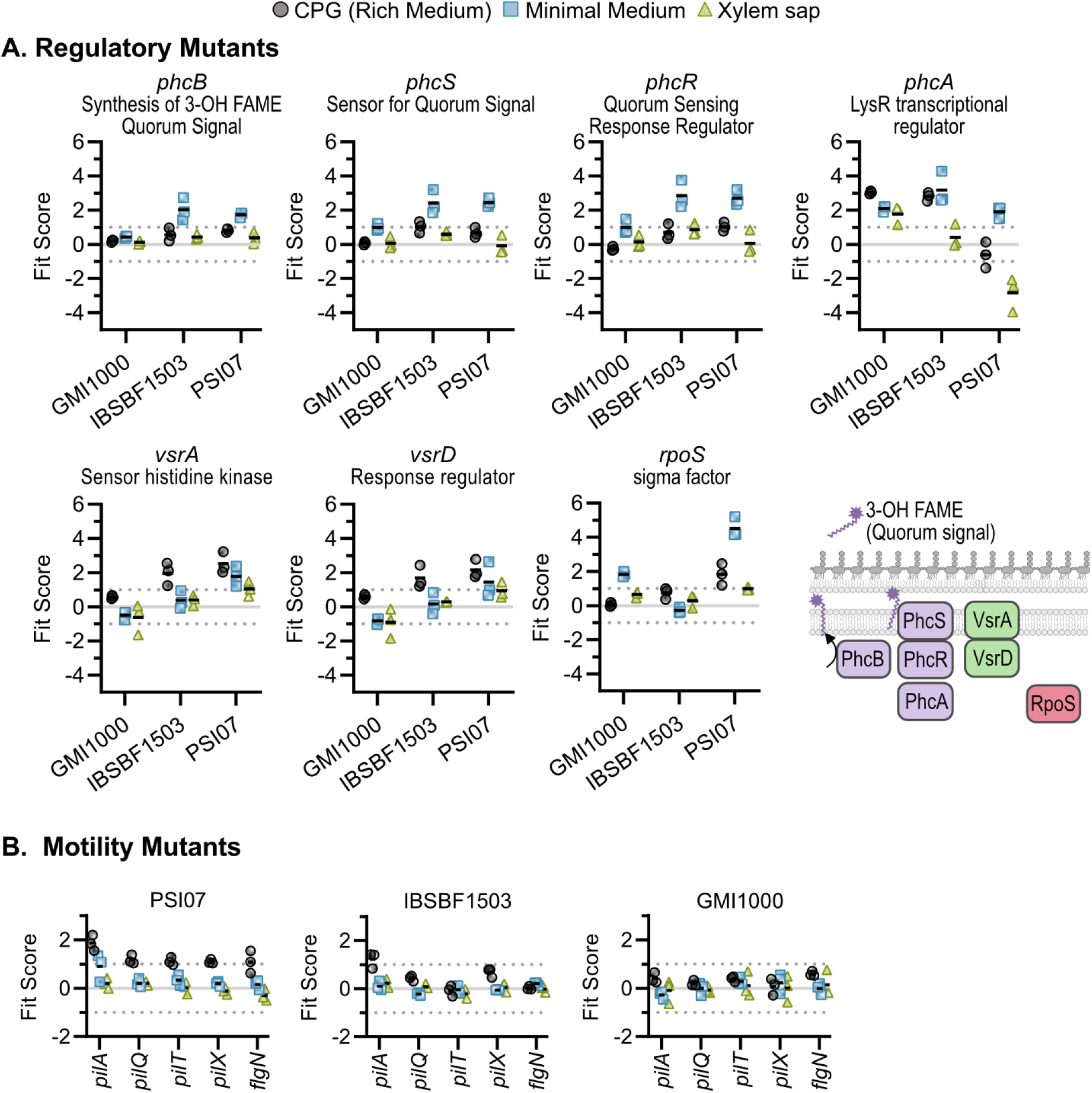
Mutants with gain-of-fitness in one-or-more strain and condition. (A) Regulatory mutants with known defects in extracellular polysaccharide (EPS) production relative to wild type: *phc* quorum sensing mutants, *vsrAD* mutants, and *rpoS* mutants. (B) Motility mutants lacking type IV pilus genes and the *flgN* flagellar chaperone genes had increased fitness in rich media in one or more strains. Graphs show Fit scores of n=3 TnSeq trials and bars show the mean. Full Fit scores and t-like test statistics are shown in Table S2.

Strain PSI07 had the most mutants with gain-of-fitness phenotypes in culture media. Four of these genes were associated with the type IV pilus, and another with a flagella-associated chaperone, FlgN (Fig 6B). Three IBSBF1503 type IV pilus mutants showed the same trend as PSI07, but with weaker phenotypes. This result suggests that these wildtype strains may express energy-consuming motility systems in liquid rich medium but not in sap or minimal medium. PSI07 had additional regulators that had gain-of-fitness phenotypes in culture media: a MarR-family transcriptional regulator nested in a hydroxycinnamic acid degradation cluster (RSp0224/RPSI07_RS00780) (13), a transcriptional regulator co-localized with an ABC transport system (RSp1579/ RPSI07_RS07405) (37), a TetR family transcriptional regulator (RSc3207/ RPSI07_RS15470), a GGDEF/EAL domain-encoding gene (RPSI07_RS05560/RSp1208), and the RelA (p)ppGpp synthase (RPSI07_RS15685/RSc1576).

## Discussion

Why might envelope genes contribute to *Ralstonia* fitness in *ex vivo* xylem sap? Recent studies have shown that clinical antibiotic use can select for pathogens that remodel their envelopes, enabling them to exclude and resist antibiotics (38). Because many envelope-associated genes were xylem sap fitness factors, we hypothesize that xylem sap contains preformed antimicrobial chemicals and proteins. Future studies should test the hypothesis that periplasmic branched oligoglucans, LPS, and other envelop genes protect *Ralstonia* from *in planta* stresses like reactive oxygen species and antimicrobial peptides (38).

There are benefits and limitations of predicting gene essentiality using transposon mutagenesis. Here we identified sets of 454-487 essential genes in three *Ralstonia* strains, of which more than half (n=244) were conserved in all three strains and predicted essential by both analysis pipelines. Unsurprisingly, many of the essential genes are involved in carbon metabolism, amino acid and cofactor biosynthesis, and central dogma processes. Identifying absolutely essential genes of human pathogens aids in antibacterial drug development research (39), but precision antimicrobial treatment for most plant pathogens is not financially viable and poses environmental concerns. Nonetheless, identifying putative essential genes has fundamental value, particularly to the *Ralstonia* research community. Creating mutants lacking these genes might be impossible or require specialized selection media as was required for the Δ*speC::*Sm mutant that had a putrescine auxotrophy (8). The essentiality of multiple amino acid biosynthesis genes in the peptone- and casamino acid-containing CPG rich medium suggests *Ralstonia* may lack effective transporters for some amino acids.

Recently, Su *et al*. predicted 464 essential genes in *Ralstonia* GMI1000 (40) using a TnSeq approach, but our libraries contain mutants with transposon insertions in 126 of these genes. Of these 126 genes, only three genes contributed to fitness more than two-fold (Fit = −1) in any of our experimental conditions, including rich medium. These discrepancies highlight the limitations of using randomly generated mutant libraries to predict gene essentiality. Targeted knockdown experiments using techniques like CRISPRi (41) could test essentiality more robustly. In a subsequent TnSeq experiment using their mutant library in living tomato plant hosts, Su *et al*. identified 131 genes as being important for *in planta* fitness (42). Of the 36 genes we identified as important for xylem sap fitness (average | Fit | > 1, | t | > 2.5 in three trials), Su *et al*. identified 16 as also being important for *in planta* fitness. This overlap suggests that *Ralstonia* experiences some of the same selective pressures in *ex vivo* xylem sap as it does in living tomato hosts, though additional genes are necessary for full fitness in the more complex *in planta* environment.

RB-TnSeq reveals inter-strain differences in fitness factors. Although model strains such as GMI1000 are well-characterized and extensively studied, there are known physiological and genetic differences between strains (1, 28, 43). To more fully capture the diversity of *Ralstonia*, we constructed barcoded transposon mutant libraries in strains from each of the three plant-pathogenic *Ralstonia* species: *R. solanacearum* (GMI1000), *R. solanacearum* (IBSBF1503), and *R. syzygii* (PSI07). While RB-TnSeq found specific metabolism- and envelope-related fitness factors in all three strains, the orthologs found to be fitness factors varied from strains to strain. For instance, GMI1000 requires *mdoGH* genes for periplasmic oligoglucan production in xylem sap, while these genes do not produce strong fitness phenotypes in the other two strains. Additionally, several amino acid biosynthesis genes were required by some by not all strains when grown in minimal media or xylem sap, including *cysU, leuB1, leuC, gltB*, and *proB* (Fig 4). These strain-to-strain differences in gene fitness profiles reveal subtle differences in the genetics that underlie the ability of all these strains to robustly grow in tomato xylem sap. Considering our results in *ex vivo* xylem sap, different genes may be important for virulence of these diverse strains in living hosts as well.

Novel environments provide strong selective pressures for bacterial evolution (44). We suggest that the number of gain-of-fitness mutations in a TnSeq screen may be an indicator of how closely the test conditions match the environment to which the bacterial isolate is adapted.

For instance, transposon insertions in over two dozen *E. coli* genes increased its growth on cheese agar (45), a foreign growth medium. Similarly, serial passage of *Ralstonia* GMI1000 in *Medicago* nodules (a foreign environment) selected for more gain-of-fitness mutations than serial passage in tomato stems (46, 47). In our study, few mutations increased fitness in sap, but many mutants increased fitness in artificial culture media, suggesting that the *Ralstonia* isolates are well-adapted to grow in tomato xylem sap. We observed that the model strain *Ralstonia* GMI1000 had fewer gain-of-function mutations in culture media than the less-studied IBSBF1503 and PSI07, suggesting that GMI1000 may be more domesticated and adapted to culture media since its isolation in 1978. Meta-analyses of TnSeq studies across multiple bacteria in novel and naturalistic conditions can test the hypothesis that gain-of-fitness phenotypes are more common when mutants grow in novel conditions.

Traditional transposon mutagenesis genetic screens pioneered molecular plant pathology and identified core virulence traits like the type III secretion system (48–50). TnSeq approaches leverage next-generation sequencing to rapidly quantify mutant phenotypes. These genome-wide fitness assays are a powerful approach to rapidly investigate basic bacterial biology and identify pathogen, commensal, and mutualist fitness factors. TnSeq studies have identified genes that promote bacterial fitness in their native environments including plant hosts (51–57) animal hosts (58–60), soft-rind cheese (45), and lake water (61). The RB-TnSeq technique that we use here is a powerful TnSeq methodology that has a low cost and technical barrier per sample, which facilitates profiling mutant fitness across in multiple strain backgrounds across many conditions (16, 62). Future studies will profile fitness of *Ralstonia* mutants *in planta* to test whether sap fitness factors also contribute to fitness in living hosts.

## Methods

### Bacterial Strains and Growth Conditions

This study uses three *Ralstonia* strains, one representative per plant pathogenic species: *R. pseudosolanacearum* GMI1000 (phylotype I sequevar 18), *R. syzygii* PSI07 (phylotype IV sequevar 10), and *R. solanacearum* IBSBF1503 (phylotype IIB sequevar 4). All isolates are pathogenic on tomato (11).

*Ralstonia* was routinely grown in CPG rich medium (per L: 1 g casamino acids, 10 g Bacto-peptone, 5 g glucose, 1 g yeast extract) at 28°C. Agar plates were supplemented with 1% tetrazolium chloride to confirm colony morphology. For minimal medium growth curves and fitness assays, strains were grown in quarter-strength M63 (per L: 3.4 g KH_2_PO_4_, 0.5 g (NH_4_)_2_SO_4_, 10 μl of 1.25 mg/ml FeSO_4_•7H_2_O, 51.7 μl of 1 M MgSO_4_; adjusted to pH 7) with 10 mM glucose.

### Construction of barcoded transposon mutant libraries

Barcoded *mariner* transposon mutant libraries were created in three wild-type strain backgrounds: GMI1000, PSI07, and IBSBF1503. The barcoded transposons were introduced via conjugation. Recipient strains (*Ralstonia* isolates) were grown overnight in 5 ml CPG at 28°C with shaking at 200 rpm. A 2 ml portion of the overnight culture was sub-cultured into 25 ml CPG and grown for 2 hours at 28°C. The donor *E. coli* strain (WM3064 background, a synthetic auxotroph requiring 300 μM diaminopimenlate [DAP]) carrying the pKMW3 (Kan^R^) barcoded *Mariner* transposon vector library (16) was thawed on ice and 1 ml was inoculated into 20 ml LB with 25 μg/ml kanamycin and 300 μM DAP and grown to mid-log phase at 37°C with shaking. Conjugations were carried out as previously described (63). Donor and recipient strains were centrifuged at 5000 x*g* for 10 min and resuspended in CPG + 300 μM DAP. Recipient strains and the donor strain were adjusted to OD_600_ of 3.0 and 1.0, respectively, and mixed in equal volumes. In total, 1.2 ml of the strain mixture was spotted onto 12 nitrocellulose filters (82 mm discs with 0.45 μM pore size cut into sixths) overlaid on four plates of CPG with 300 μM DAP and without glucose. Mating was allowed to occur for 3-4 hours at 28°C. Filters were pooled in 20 ml CPG broth and vortexed in a 50 ml tube for 30 s to dislodge bacteria. The suspension was adjusted to 80 ml and evenly spread over 400 CPG plates (200 μl per plate) with 25 µg/ml kanamycin. To calculate transformation efficiencies, a portion of each suspension was dilution plated on CPG with kanamycin (to quantify transformants) and without kanamycin (to quantify total *Ralstonia* cells). Transformation efficiencies were: GMI1000, 3.9 × 10^−5^; PSI07, 8.6 × 10^−3^; IBSBF1503, 1.5 × 10^−4^. Selection plates were incubated for 2 days at 28°C. Cells were harvested by scraping and pooling approximately 2 × 10^6^ colonies in CPG broth with kanamycin. Density was adjusted to OD_600_ = 0.25 in 200 ml and grown at 28°C with shaking until OD_600_ reached 1.0. Individual 1 ml aliquots were preserved in 20% v/v glycerol (final concentration) at −80°C.

### Mapping transposon insertion sites

We used a previously described protocol for mapping the transposon insertion locations and to link these mutants to their unique DNA barcode sequences (16). Full details describing this protocol are available (16). Briefly, genomic DNA was extracted using the Qiagen DNeasy Blood and Tissue Kit, with RNase A treatment, according to the manufacturer’s recommendations. The DNA was sheared to ~300 bp using a Covaris S220, and the DNA was repaired and A-tailed using the NEBNext DNA Library preparation kit for Illumina (New England Biolabs). After ligating Illumina compatible adapters to the DNA fragments, we PCR amplified the transposon insertion junctions using primers with Illumina P5 and P7 sequences. The final PCR product was purified using AMPure XP beads and assessed for quantity and size using an Agilent Bioanalyzer DNA1000 chip. Samples were sequenced on a HiSeq2500 instrument at the QB3 Berkeley Genomics Center using 2 × 150 reads. TnSeq reads were analyzed with a custom Perl script, MapTnSeq.pl, which assigns each unique barcode sequence to a corresponding location in the genome. Each barcode was mapped to a single insertion site by identifying barcodes that consistently map to a unique location in the genome using DesignRandomPool.pl. All scripts used are available at https://bitbucket.org/berkeleylab/feba/src/master/bin/ (16, 62).

### Essential gene calculations

Based on the TnSeq data, we used standard computational methods (62) to predict which genes are likely essential for growth in CPG. Briefly, this analysis predicts gene essentiality based on genes that lacked transposon insertions in the central 80% of the coding region. We excluded two categories of genes from the essentiality calculation. Genes sharing high nucleotide identity with other genes in the genome (measured via BLAT) (64) were excluded because we could not map transposon insertions to highly similar genes. Second, we excluded genes below a minimum length because they are less likely to be disrupted. Briefly, this cutoff is determined using our previously described analysis pipeline (18) using saturation of *mariner* transposon insertions to calculate the length at which genes have a 1% chance of being missed due to random chance. Due to differences in insertion mutant coverage depth between the transposon libraries, the minimum gene length considered was 450 bp for GMI1000, 325 bp for PSI07, and 400 bp for IBSBF1503. For comparison, we also used the Bio-Tradis pipeline (github.com/sanger-pathogens/Bio-Tradis) (17) to predict essential genes in these strains, using the same minimum gene length cutoffs and excluding insertions in the first and last 10% of the coding region from analysis.

### Orthologous gene predictions

Orthologous genes were predicted using the “Genome Gene Best Homologs” tool from the JGI IMG database (19). We searched for homologs in PSI07 and IBSBF1503 against GMI1000 as the reference genome using the JGI IMG default identity threshold of 60%. The results, originally reported with IMG locus tags, were matched to the corresponding NCBI locus tags by gene sequence.

### Tomato xylem sap harvesting

Xylem sap was harvested from susceptible tomato plants (cv. Moneymaker and Bonny Best). Most experiments were performed with sap from Moneymaker tomato plants grown in the Oxford Tract Greenhouse in Berkeley, CA. Bacterial growth curves were independently replicated with sap from tomato plants grown in a growth chamber (28°C with a 12 hr day/night cycle) at UW-Madison.

Xylem sap was harvested by de-topping each plant at the cotyledon juncture with a razor blade and allowing sap to pool on the stump (65). To reduce cytoplasmic contamination, the sap that accumulated in the first few minutes was discarded; the stump was washed with diH_2_O and gently blotted dry. Sap was then pipetted and pooled into a 50 ml conical over a collection period of 3 hours. Pooled sap was centrifuged at 5,000 *xg* RT for 10 minutes and the supernatant was filter sterilized using filters with 0.22 μm pores (Thermo Scientific #725-2520). Sap was aliquoted and stored at −20°C until use. Each batched pool of xylem sap was collected from approximately 70 plants.

### *R. solanacearum* growth in tomato xylem sap

Colonies of each strain were inoculated into 6 ml of CPG and grown overnight at 28°C with shaking at 200 rpm for a total of three biological replicates. Cells from stationary phase cultures were pelleted at 13,000 *xg* and washed twice in 1 ml of ddH_2_O. The washed pellet was resuspended in ddH_2_O and adjusted to a final OD_600_ of 0.02. Each growth condition was inoculated with 7 μl of each culture at a starting cell density of ~10^5^ CFU/ml in 48-well plates (Corning #353078), for a total of two technical replicates per biological replicate. Growth was measured by dilution plating at 0-, 4-, 8-, 24-, 32-, and 48-hour time points. In parallel, measurements for 12-, 16-, and 20-hour time points were taken from independent cultures that were started 12 hours after the first, using the same cultures, growth media, and technique described above.

### Fitness experiments with transposon libraries

For genome wide fitness experiments, we adapted established methods (16). Per condition (minimal medium, rich medium, or Moneymaker tomato xylem sap), three 1 ml aliquots of each transposon library (GMI1000, PSI07, and IBSBF1503) were thawed on ice and revived in separate 100 ml flasks of CPG with 12.5 mg/ml Kanamycin with shaking incubation at 28°C for 16-20 hours. Once cultures reached an OD_600_ of 0.2 to 0.5, cells were pelleted by centrifuging for 10 min at 5000 *xg* RT and resuspended in 2 ml of rich or minimal medium or sterile H_2_O (xylem sap experiments). Cells were washed 3 times in media or xylem sap. An aliquot of the washed cells (>10^9^ cells) was retained (pelleted and frozen) for a “time 0” control. For rich and minimal medium experiments, cells were seeded into 5 ml at OD_600_ = 0.02 (~2-5×10^7^ total cells). Rich medium cultures were grown to saturation (20-24 hours; 5-6 cell doublings) and minimal medium cultures were grown for over 48 hours (5-6 cell doublings for PSI07 and GMI1000 and 4 doublings for IBSBF1503). For xylem sap experiments, cells were resuspended in 20 ml of xylem sap at a starting cell density of 10^6^ cells/ml (2 × 10^7^ cells total) and grown for 25 hours (7-8 doublings for GMI1000 and 8-9 doublings for PSI07). Genomic DNA was extracted (Qiagen DNeasy Blood and Tissue Kit) from the “time 0” and cells harvested after growing in differential media.

### BarSeq and Fitness Calculations

Prior to PCR, the quality if total genomic DNA isolated from fitness experiments was assessed on an agarose gel, while DNA concentration was determined by NanoDrop. Barcodes were PCR amplified from each sample using the previously described reaction protocol (16). For most experiments, we used indexed P2 oligos and a mix of non-indexed P1 oligos of variable length (2-5 bases) to stagger the reads. For some experiments, we used indexed versions of both the P1 and P2 oligos to minimize incorrectly assigned indexes in Illumina HiSeq4000 runs. Following PCR, amplicon mixtures from each sample were pooled in equal volumes, purified over a Zymo DNA Clean and Concentrator column, and eluted with water. Prior to sequencing, the quality of cleaned and concentrated amplicon pools was assessed using a Bioanalyzer. qPCR was used to determine how much of each sample to load for sequencing. Barcode amplicons were sequenced on a HiSeq4000 at QB3 Berkeley Genomics Center with 96 samples multiplexed (50 base pair reads, single end).

Sequencing data was analyzed using the BarSeq pipeline, available at https://bitbucket.org/berkeleylab/feba/src/master/bin/. For each competitive fitness assay, fitness score (“Fit scores”) for each gene were calculated as the log_2_ ratio between barcode abundance after outgrowth in a condition vs. its abundance in the time 0 sample. Each Fit score is the weighted average of fitness values for all mutant strains with transposon insertions in a given gene. These Fit scores are normalized across the genome such that a gene with neither a fitness cost nor benefit has a value of 0. Significance was determined based on an absolute t-like test statistic with a threshold of | t | > 2.5 in all three RB-TnSeq trials.. The t-like test statistic considers the consistency of the fitness scores for all barcoded mutants for each gene in the experiment as previously described in detail (16).

### Data availability

The raw reads used for TnSeq mapping is available in the NCBI SRA under accession PRJNA629015. The fitness browser (http://fit.genomics.lbl.gov) offers a graphical user interface for exploring the fitness data from these experiments, orthology of the genes between these strains and other Gram negative bacteria, and cross-references to Kegg, Paperblast, and NCBI databases.

## Acknowledgements

We thank Steve Lindow, Caitilyn Allen, and Jeff Flynn for useful discussions. This work used the Vincent J. Coates Genomics Sequencing Laboratory at UC Berkeley, supported by NIH S10 OD018174 Instrumentation Grant. Oxford Tract Greenhouse Staff and the QB3 Berkeley Genomics Center provided technical assistance.

This work was supported by USDA NIFA #2018-67012-31497 awarded to TLP; partial support was provided by a UC Berkeley SURF Rose Hills Independent Fellowship awarded to KES. The funders had no role in study design, data collection and interpretation, or the decision to submit the work for publication.

Mention of trade names or commercial products in this publication is solely for the purpose of providing specific information and does not imply recommendation or endorsement by the U.S. Department of Agriculture. USDA is an equal opportunity provider and employer.

## Supplementary Data

**Fig S1.**
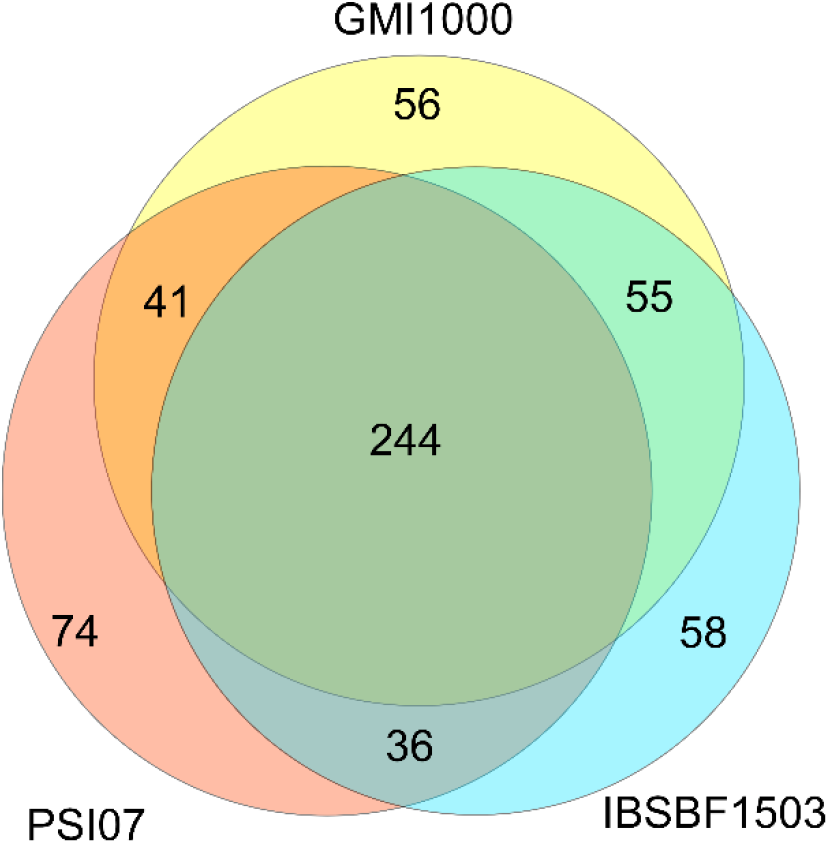
Venn diagram of putative essential genes in *Ralstonia* strains GMI1000, IBSBF1503, and PSI07. Orthologous predicted essential genes are represented proportionally in Venn diagram intersections. Genes were included in the Venn diagram only if they were predicted to be essential by both analysis pipelines used in this study (17, 18).

**Fig S2.**
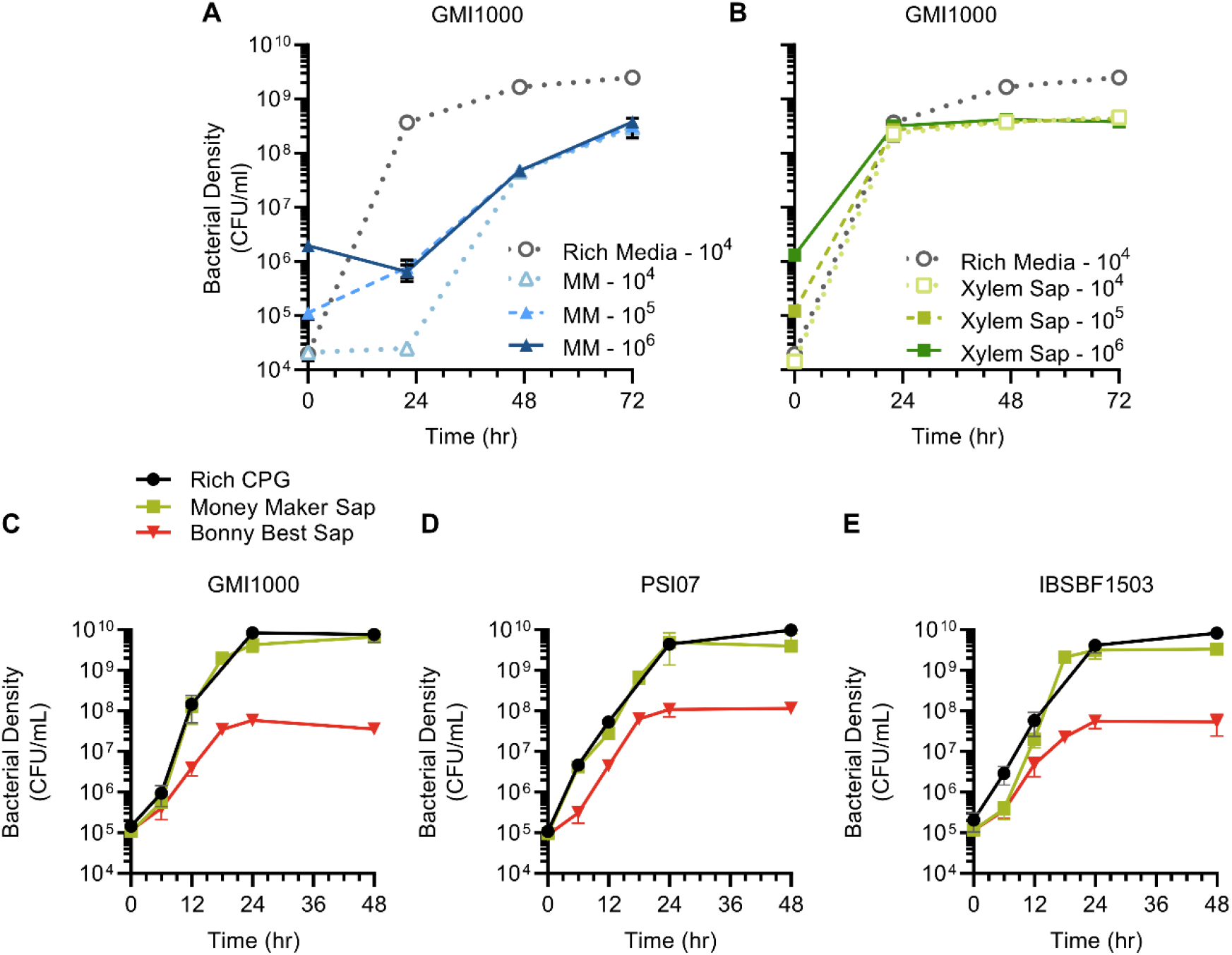
Growth of *Ralstonia* in rich medium, minimal medium, and xylem sap from multiple susceptible tomato cultivars. (A-B) To determine the carrying capacity of sap and culture media, *Ralstonia* strain GMI1000 was inoculated into rich CPG medium, quarter strength M63 medium, or xylem sap from Moneymaker tomato at 10^4^, 10^5^, or 10^6^ CFU/ml starting density. (C-E). Wildtype *Ralstonia* strains GMI1000, PSI07, and IBSBF1503 were grown in sap from Moneymaker and Bonny Best tomato. All strains were inoculated in biological triplicate into three pools of xylem sap incubated at 28°C with shaking. Cell density was measured by dilution plating.

**Table S1: Gene essentiality predictions for GMI1000, PSI07, and IBSBF1503**

**Table S2: Genome-wide Fit scores for GMI1000, PSI07, and IBSBF1503**

